# Antagonism of stress granules key for SARS-CoV-2 infection and pathogenesis

**DOI:** 10.64898/2026.06.16.732644

**Authors:** R. Elias Alvarado, Jennifer Chen, Kumari G. Lokugamage, Yiyang Zhou, Leah K. Estes, Angelica Morgan, Yani Ahearn, Xiangxue Deng, Lilin Lai, Arian Moayyed, William Meyers, Jessica A. Plante, Ken S. Plante, David H. Walker, Xuping Xie, Mehul S. Suthar, Bryan A. Johnson, Vineet D. Menachery

## Abstract

Viruses must subvert host responses to facilitate successful infection. While most antiviral responses are associated with interferons, stress granules (SG) are another barrier to viral infection by inducing translation arrest. To combat SG activity, viruses have evolved mechanisms to disrupt formation and disassemble these complexes. Our prior studies identified residues in SARS-CoV-2 NSP3 (Y138/F145) and nucleocapsid (F17) that independently antagonize SG activity. Disrupting these key residues in NSP3 or nucleocapsid attenuated viral replication, but only modestly impacted *in vivo* pathogenesis suggesting overlap in SG antagonism partially compensate for the individual losses. In this study, we evaluated a SARS-CoV-2 mutant (YF/F17A) that combines the NSP3 and N mutations. We find that loss of both SG antagonizing functions attenuates SARS-CoV-2 replication *in vitro*. While no changes are seen in type I IFN sensitivity, attenuation corresponds to increased induction of SGs. Importantly, the SARS-CoV-2 YF/F17A mutant has significant attenuation *in vivo* with reduced viral replication, less weight loss, and limited immune pathology. Notably, infection with the YF/F17A mutant stimulated less interferon and inflammation. Despite these muted host responses, the YF/F17A mutant stimulated robust protection against subsequent challenge with WT SARS-CoV-2. Overall, the study highlights the importance of SG control for SARS-CoV-2 infection and offers a novel, interferon independent target for vaccination and therapeutic treatment going forward.

**Importance:** This study demonstrates that SARS-CoV-2 uses multiple mechanisms to block host stress granules during infection. Knocking out both N and NSP3 mediated antagonism of stress granules attenuates viral replication and disease caused by SARS-CoV-2. Importantly, while most therapeutics target key viral processes or induce interferon pathways, this study shows stress granule activation as a novel approach to attenuate and treat coronavirus infection.

## Introduction

Coronaviruses are positive-sense, enveloped RNA viruses that have posed significant global threats to human health and led to three pandemics this century (1–3). To establish infection, viruses must interact with host cellular components in order to establish infection and generate progeny virus (4). Therefore, understanding these host-virus interactions is paramount for developing new antivirals and vaccine candidates.

Eukaryotic cells have evolved defense mechanisms to sense and respond to cellular stresses, including viral infections (5–7). When encountering stress, cells rapidly arrest translation by activating stress responsive kinases that disrupt the formation of the eukaryotic translation initiation factor 4F (eIF4F) complex (8). Accumulated untranslated mRNAs engage with host proteins to form cytosolic foci termed stress granules (SGs). SGs are transient ribonucleoprotein (RNP) condensates formed via liquid-liquid phase separation (LLPS), in which stalled translation preinitiation complexes, cellular mRNAs, small ribosomal subunits, and RNA-binding proteins such as Ras GTPase-activation protein-binding 1 (G3BP1) and Fragile X mental retardation proteins (FMRPs) coalesce into a concentrated phase (9). Host cells respond to stress such as viral infections by forming SGs that can further disrupt host translation by sequestering these key factors. To counteract SGs, some viruses respond by hijacking, delaying, or disrupting SG formation during infection to ensure host machinery maintains translation of viral proteins (10–12). Taken together, SGs are a critical obstacle for viruses to overcome during infection.

Prior studies have shown that SARS-CoV-2 inhibits SG formation though multiple virus-host protein interactions (13–20). The nucleocapsid (N) protein of SARS-CoV-2 is a key structural protein that has several roles during infection and is primarily known for binding viral RNA and facilitating its incorporation into new virions (21). N also suppresses innate immune responses and supports viral RNA replication and transcription (22–24). These different roles for N can be regulated by kinetics, viral RNA regulation, and protein phosphorylation (25–27). Recently, work from our lab and others has shown that residue F17 in N protein binds to the G3BP1 NTF2L domain with high affinity and promotes SG disassembly (13, 28, 29). Interestingly, we found that mutating residue F17 to alanine (F17A) in a SARS-CoV-2 mutant resulted in persistent SGs at late time points (24 hours post infection (HPI) (13). These observations indicate that N is a strong antagonist of SGs by inhibiting their functions at late times during infection.

In addition to N, SARS-CoV-2 nonstructural protein 3 (NSP3) can also inhibit SG formation. Our group identified that the hyper-variable region (HVR) of NSP3 binds the SG-related Fragile X mental retardation protein 1 (FXR1) via interaction with ubiquitin associated protein 2 like (UBA2PL) (30). Disruption of the HVR region via alanine scanning, resulted in enhanced SG formation and attenuated viral infection (6 HPI)(30). We subsequently identified two key aromatic residues in the NSP3 HVR, tyrosine (Y) 138 and phenylalanine (F) 145 responsible for binding UBA2PL (14). A SARS-CoV-2 mutant disrupting these residues (YF) had SG formation at early times (6 HPI) that was subsequently inhibited at later stages of infection (10 HPI). The disruption of SGs at late times corresponds to increased N levels at late times during infection. Together, our results from two separate studies demonstrate that NSP3 and N selectively target SGs at different times during infection (14). With such a significant investment in genetic capital, these results suggest control of SG formation is paramount to SARS-CoV-2 infection and pathogenesis.

To explore this idea, we examined the impact of SG activation by generating recombinant SARS-CoV-2 bearing mutations in the NSP3-HVR and N (YF/F17A) that prevent binding to FXR1 and G3BP1, respectively. We demonstrate that SARS-CoV-2 YF/F17A mutant has reduced viral replication *in vitro* which corresponded to increased SG formation rather than type I IFN activation. We also observed attenuated pathogenesis *in vivo* with reduced disease and host responses. Notably, we show that despite diminished immune activation during primary infection, the YF/F17A mutant conferred protective immunity following vaccination and subsequent rechallenge with WT SARS-CoV-2. Overall, our studies demonstrate the importance of controlling SGs for SARS-CoV-2 infection and pathogenesis. Similarly, these findings demonstrate SG activation as a viable strategy for vaccination and therapeutic development for coronavirus infection.

## Results

### Disruption of NSP3 and N stress granule antagonism attenuates SARS-CoV-2 infection

Our prior work identified two distinct mechanisms that disrupt SG formation during SARS-CoV-2 infection: 1) F17A mutation in SARS-CoV-2 N, that disrupts binding to G3BP1/2 (13), and 2) Y138A/F145A (YF) in SARS-CoV-2 NSP3, disrupts binding to FMRPs (14, 30). Although both mutants are moderately attenuated, their targeting of distinct SG components and differential effects on the temporal kinetics indicate that they disrupt two independent mechanisms of SG activation (**Fig. 1A**) (13, 14, 30). Consequently, we concluded that mutation at both interfaces may fully restore SGs during SARS-CoV-2 infection. Using our reverse genetics system (31, 32), we generated an NSP3/N double mutant (YF/F17A) in the SARS-CoV-2 WA-1 strain backbone(33, 34) (**Fig. 1B**).

**Figure 1:**
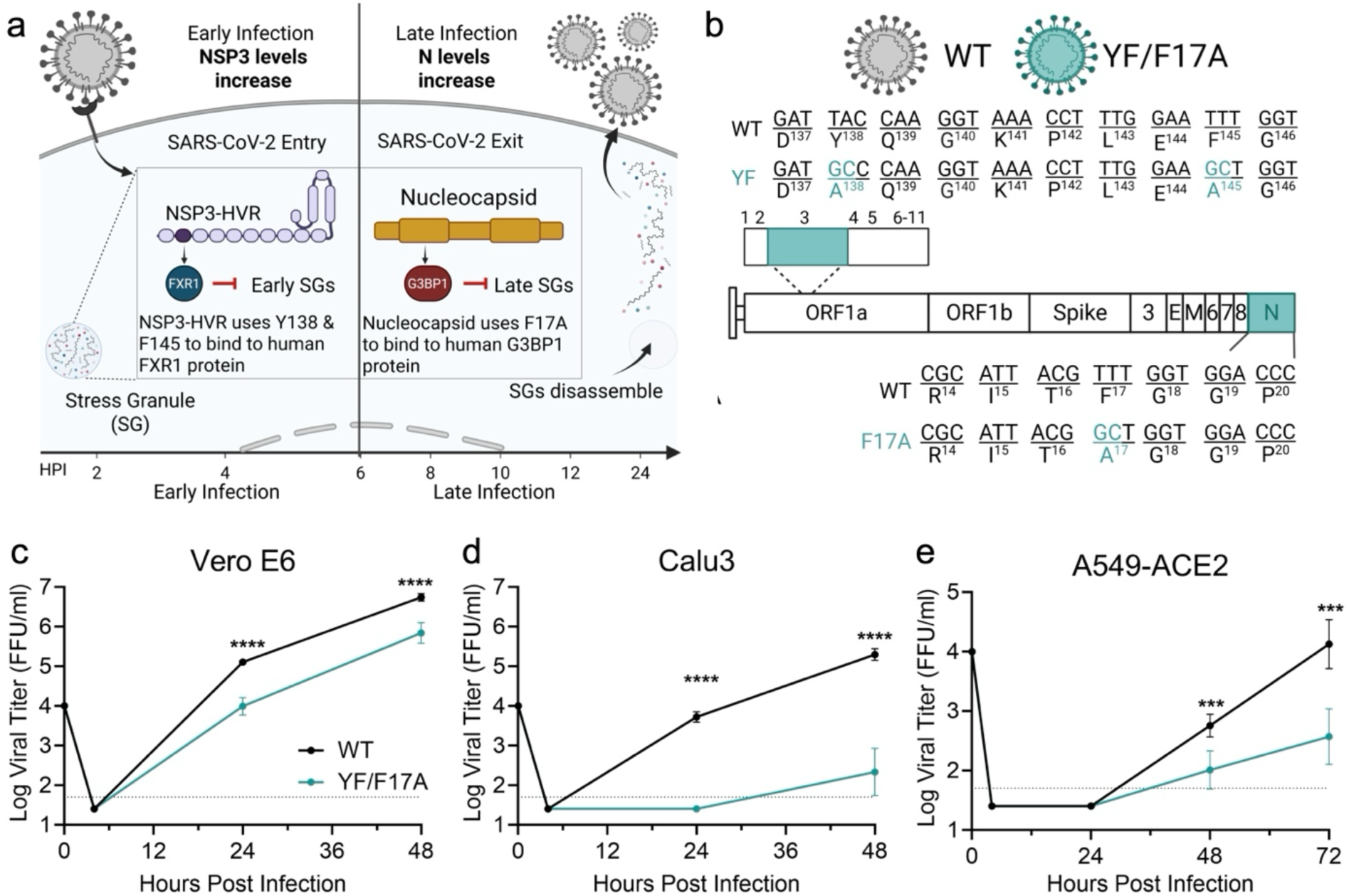
SARS-CoV-2 YF/F17A mutant has attenuated viral replication *in vitro*. (**a**) Proposed model of SARS-CoV-2 SG regulation by NSP3 and N protein at early and late stages of infection, respectively. (**b**) Schematic of SARS-CoV-2 double mutant (YF/F17A) with NSP3-YF and N-F17 mutations to alanine (teal) introduced in Washington 1(WA-1) strain. (**c-e**) Viral titers in Vero E6 (**c**), Calu3 (**d**), or A549-ACE2 (**e**) cells infected with WT (black) or YF/F17A (teal) SARS-CoV-2 at an MOI of 0.01 (n = 6 from two experiments, each with three biological replicates). Data are representative of mean ± SD. Statistical analysis measured by two-tailed Student’s t test. *P ≤ 0.05; **P ≤ 0.01; ***P ≤ 0.001, ****p<0.0001. Figure 4.1A-B were created with Biorender.com.

We first evaluated the viral replication of YF/F17A compared to WT SARS-CoV-2 in cell culture using a low multiplicity of infection (MOI 0.01). Three cell models were selected for this analysis: Vero E6, an interferon (IFN) deficient cell line (35, 36); Calu3, a human bronchial cell line with intact interferon IFN responses (34, 37, 38); and A549-ACE2 cells, a lung adenocarcinoma cell line expressing human ACE2, with intact IFN responses and known stable SG activation (37, 39). Vero E6 growth curves revealed significant attenuation between the WT and YF/F17A at 24- and 48-hours post infection (HPI) (**Fig. 1C**). In Calu3 cells, YF/F17A mutant virus was not detected at 24 HPI for YF/F17A mutant, while WT SARS-CoV-2 grew to 10^4^ FFU/ml (**Fig. 1D**); by 48 HPI, YF/F17A was observed, but was attenuated 3-log compared to control (**Fig.1D**). In A549-ACE2 cells, the YF/F17A mutant replication was similarly attenuated relative to WT SARS-CoV-2 at 48- and 72 HPI. (**Fig.1E**). Collectively, these data show that the loss of SG antagonism via the mutation of both the NSP3-FXR1 and N-G3BP1/2 interactions mechanisms strongly attenuates SARS-CoV-2 replication *in vitro*.

### YF/F17 mutant attenuation not primarily due to type I interferon responses

Both NSP3 and N are multi-functional proteins in the context of CoV infection with roles in viral RNA and protein regulation, host-virus interactions, and modulation of host immune responses (40). Given the level of attenuation of the YF/F17A mutant, the combined disruption may unexpectedly impact type I IFN responses. While neither the YF or F17A individual mutants induces significant changes in innate immune responses (13, 14, 30), we evaluated YF/F17A replication for IFN sensitivity on Vero E6 cells which lack capacity to produce type I IFN, but respond to exogenous treatment (36). Pretreatment with 100 U of recombinant type I IFN (IFN-α) prior to infection, WT SARS-CoV-2 displayed modest sensitivity to type I IFN pretreatment with ∼1-log reduction in viral titers at 24 HPI. (**Fig. 2A**). Similarly, the YF/F17A mutant also displayed only modest IFN sensitivity with a ∼1 log reduction in viral titer at 24 HPI (**Fig. 2A**). Unlike IFN sensitive SARS-CoV-2 mutants that often display >3-log attenuation after pretreatment (41), these data are consistent with our prior studies with the YF and F17A individual mutants and argue that type I IFN signaling is not the predominant cause of the YF/F17A mutant attenuation (14).

**Figure 2:**
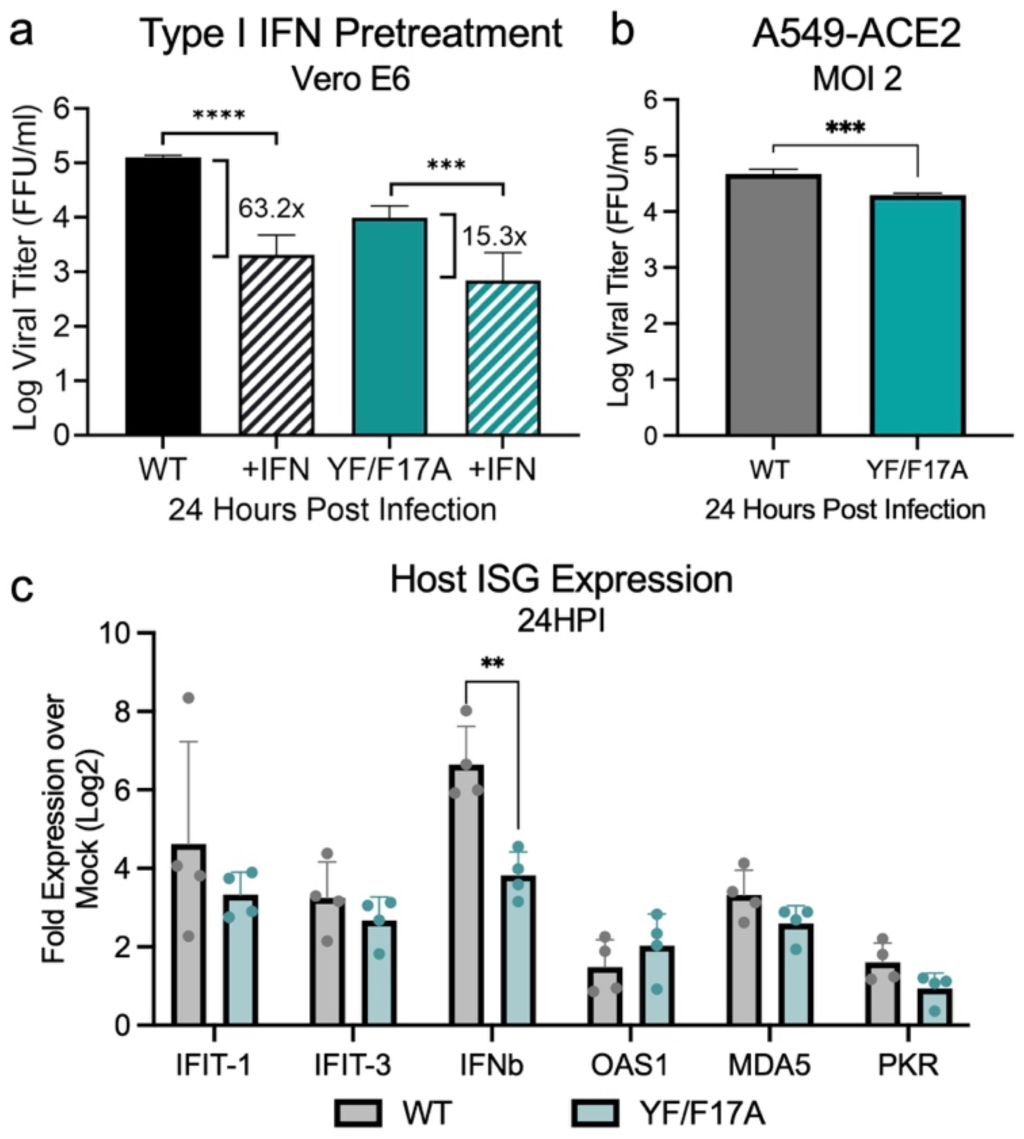
Decreased YF/F17A mutant replication is not driven by innate immune responses. (**a**) Vero E6 cells were pretreated with 100 U of recombinant type 1 IFN (hashed bar) or mock (solid bar) for 16 hours prior to infection. Cells were subsequently infected with either SARS-CoV-2 WT (black) or YF/F17A mutant (teal) at a MOI of 0.01. Viral titers were measured at 24 HPI. The fold change relative to mock control is shown. (**b -c**) A549-ACE2 cells were mock, WT, or YF/F17A infected at an MOI 1 for 24 hours with viral titer shown at endpoint. (**c**) At 24 HPI, total RNA was collected, and host mRNA expression was quantified by RT-qPCR. C_t_ values were normalized to human GAPDH and expressed as fold change (log_2_) of expression of indicated genes. Each dot represents a replicate, with the means for each target displayed, ± SD (error bars).

While the YF/F17A mutant is not more sensitive to type I IFN pretreatment, it is possible that infection may induce augmented type I IFN responses resulting in reduced capacity to infect bystander cells. To explore this possibility, we infected human A549 cells expressing hACE2 with WT and YF/F17A at a high MOI (2). Following infection, we found both WT and YF/F17A had similar titers 24 HPI (**Fig. 2B**). Examining host RNA expression, we found few changes in a panel of type I interferon (*IFNB1)* and interferon stimulated genes (*IFIT1, IFIT3, OAS1, MDA5, PKR*), consistent with previous work (**Fig. 2C**) (14). Most ISGs maintained similar levels of expression between WT and YF/F17A mutant infected cells. However, IFNβ levels were augmented in WT as compared to the YF/F17A mutant. Together, the ISG and IFN gene expression data indicate that the YF/F17A mutant does not increase host ISG responses. Overall, these data are consistent with prior studies and confirm that attenuation of the YF/F17A mutant is not due to changes on increases in host innate immune responses.

### Reduced YF/F17 mutant replication corresponds to high levels of stress granule formation

Our prior studies have indicated that loss of either NSP3 interaction with FMRP proteins or N interaction with G3BP1 increases SG activation (13, 30). Here, we sought to evaluate induction of SGs in the YF/F17A mutant following infection of A549-ACE2 cells. Briefly, A549-ACE2 cells were plated and infected with WT SARS-CoV-2, YF/F17A mutant or mock infected. At 6, 10, and 24 HPI, cells were fixed, permeabilized, stained, and evaluated for SG formation by examining number of cells staining positive for both SARS-CoV-2 N (red) and host G3BP1 (green) foci. Consistent with prior studies (13, 14, 30), WT SARS-CoV-2 fails to induce SGs at any time point following infection while the YF/F17A mutant has strong induction comparatively (**Fig. 3**). At 6 HPI, SG formation is not observed in mock, WT, or YF/F17A mutant (**Fig. 3A-D**). While N protein should be produced by this timepoint, very few cells in either infected group are positive suggesting limitations on staining sensitivity. By 10 HPI, WT SARS-CoV-2 still has minimal SG formation, similar to mock, as measured by G3BP1 foci, but now accompanied by significant N staining (**Fig. 3E-F**). In contrast, we observed a major increase in cells with both G3BP1 foci and N staining in the YF/F17A mutant (**Fig. 3G-H**) indicating the loss of SG control. The loss of SG control in the YF/F17A mutant is maintained at 24 HPI, as over 1/4 of cells show evidence of SG induction compared to <3% in the WT infected group (**Fig. 3I-L**). Together, these results demonstrate that loss of both NSP3 and N antagonism allows viral infection to induce a significant level of SGs.

**Figure 3:**
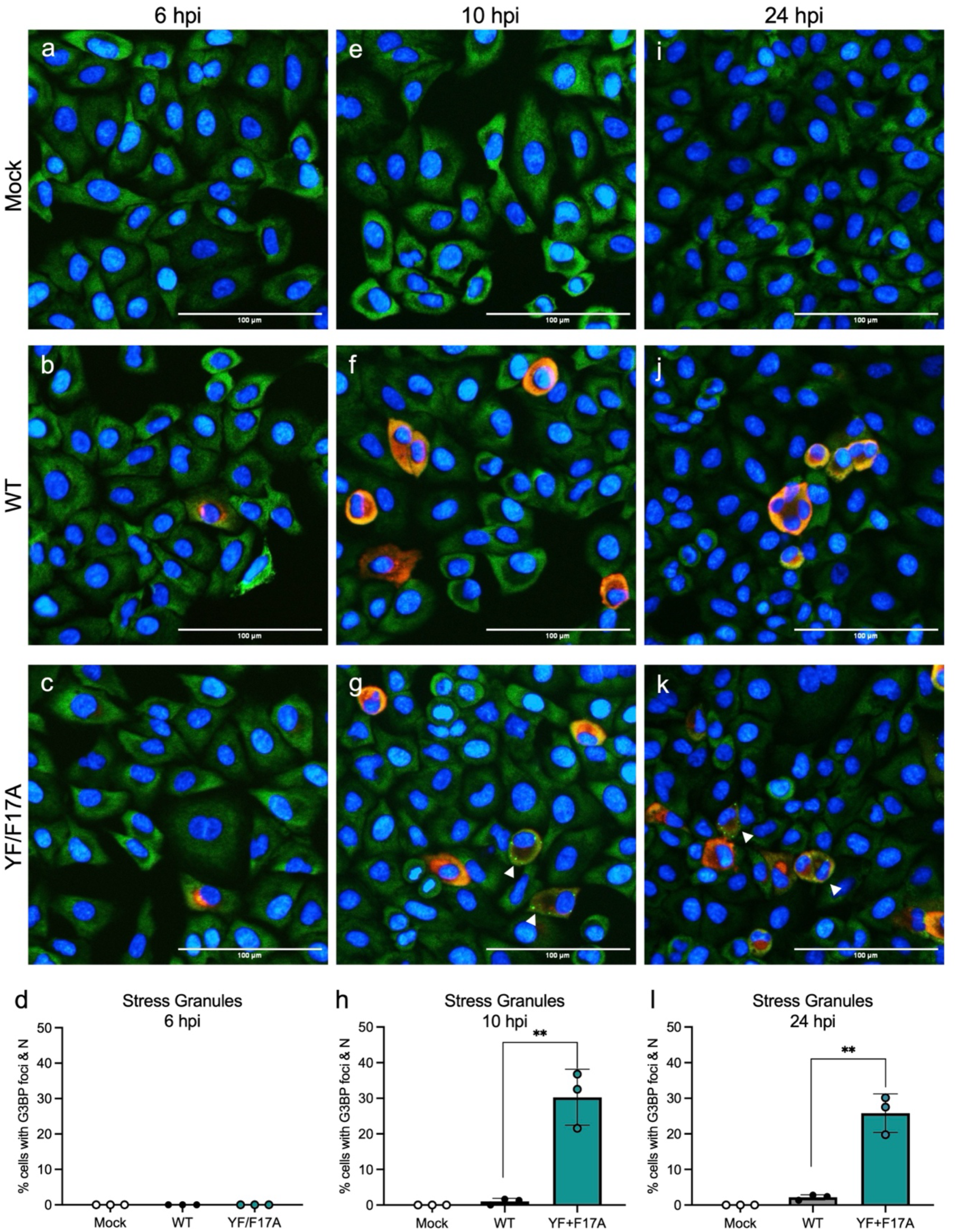
Stress granule formation is observed in YF/F17A mutant. (a-l) A549-ACE2 cells were infected with mock PBS, WT or YF/F17A mutant at MOI 2 and visualized and quantified for SG formation a (**a-d**) 6 HPI, (**e-h)** 10 HPI, and (**i-l**) 24 HPI. G3BP1 (green), Nucleocapsid (red), and nuclei (blue) were labeled to visualize by immunofluorescence imaging. The total percent of cells with G3BP1+ foci at 6, 10, and 24 HPI were calculated. All quantitative data were calculated using Fiji (ImageJ) software and are shown as mean ± SD. Statistical analysis measured by two-tailed Student’s t test. **P ≤ 0.01.

### YF/F17 mutant is unable to reverse arsenite induced SGs

To further evaluate SG activation and control, we examined infection after sodium arsenite treatment, a known inducer of SGs (42). Briefly, A549-ACE2 cells were infected with mock (PBS), SARS-CoV-2 WT or YF/F17A mutant as described above; 30 minutes prior to infection timepoint, sodium arsenite containing media was used to replace culture media followed by inactivation and staining (**Fig. 4**). Consistent with prior studies, treatment with arsenite induce formation of SGs in mock infected cells at 10 HPI as indicated by G3BP1 foci (**Fig. 4A**). For evaluating SG formation during infection, we focused on cells that expressed both G3BP1 foci (green puncta) and N (red). Examining WT SARS-CoV-2 at 10 HPI, we observed that cells with N staining lacked G3BP1 foci observed in non-infected bystander cells (**Fig. 4B**). In contrast, the YF/F17A mutant had many cells that were double positive for N and G3BP1 foci (**Fig. 4C**). Examining 24 HPI, we observed a similar trend with mock infected and bystander cells showing evidence of SG formation by G3BP1 foci (**Fig. 4D-F**). In WT infection at 24 HPI, cells expressing N had very low staining with G3BP1 foci and had evidence of cell fusion (**Fig. 4E)**. In contrast, N and G3BP1 foci double positive staining remained high in the YF/F17A mutant (**Fig. 4F**). We observed a significant increase in SG formation in the YF/F17A mutant compared to WT at both 10 and 24 HPI (**Fig. 4G & H**). We also noted that while N positive cell numbers were equal at 10 HPI, WT infected cells were 2-fold more abundant than YF/F17A mutant at 24 HPI with or without arsenite treatment (**Fig. 4I&J**). Together, the results show that the YF/F17A mutant is unable to limit externally induced SG formation and has reduced infection over time.

**Figure 4:**
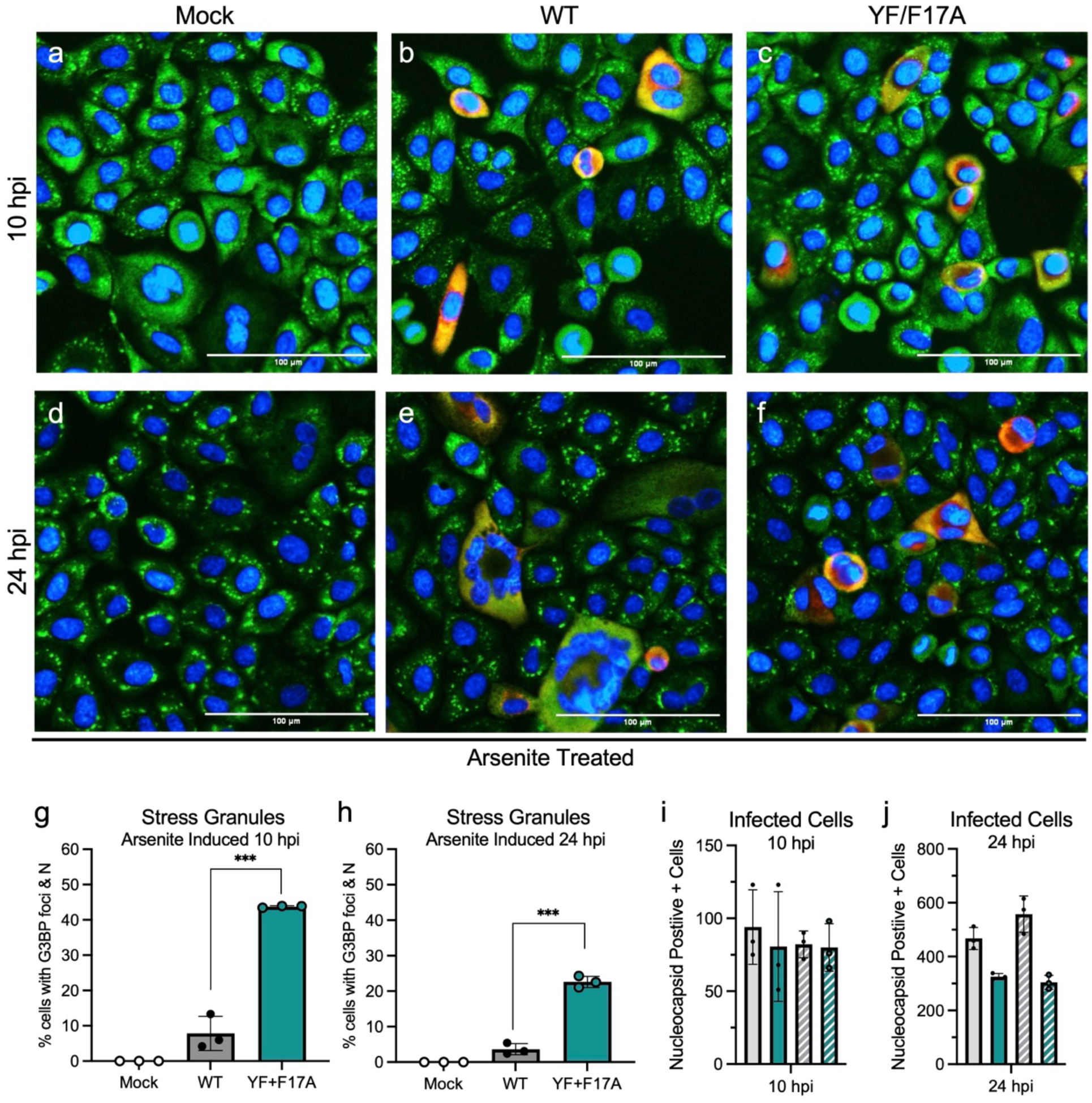
YF/F17A mutant is unable to control arsenite induced stress granule formation (a-f) A549-ACE2 cells were infected with mock PBS, WT or YF/F17A mutant at MOI 2 and visualized for SG formation at (**a-c**) 10 HPI and (**d-f**) 24 HPI. G3BP1 (green), Nucleocapsid (red), and nuclei (blue) were labeled to visualize by immunofluorescence imaging. (**g-h**) The total percent of cells with G3BP1+ foci at (**g**) 10 HPI and (**h**) 24 HPI were calculated. (**i-j**) Total number of N positive cells was calculated for WT (Grey) and YF/F17A mutant (teal) following control (solid) and arsenite treatment (hashed) at (**i**) 10 HPI and (**j**) 24 HPI. All quantitative imaging data were calculated using Fiji (ImageJ) software and are shown as mean ± SD. Statistical analysis measured by two-tailed Student’s t test. **P ≤ 0.01.

### YF/F17A mutant significantly attenuated weight loss and viral replication *in vivo*

Having established significant *in vitro* attenuation and robust induction of SGs, we next sought to evaluate the YF/F17A mutant *in vivo*. Using the Syrian golden hamster model for SARS-CoV-2 infection (43–46), we intranasally challenged animals with 10^5^ FFU of either WT SARS-CoV-2, YF/F17A mutant, or PBS (mock) and monitored daily for weight loss and clinical signs of disease (**Fig. 5A**). At 2-,4-, and 7- days post infection (DPI), nasal washes were collected, followed by euthanasia and lung tissue harvested for measurement of viral titer, RNA, and histopathology. Consistent with prior studies, hamsters infected with SARS-CoV-2 WT had weight loss beginning at 2 DPI and continues until it peaked at ∼15% at 6 DPI (**Fig. 5B**). In contrast, YF/F17A infected hamsters were significantly attenuated in weight loss compared to WT infection throughout the time course. While mock infected animals gained weight, the YF/F17A mutant infected hamsters maintained their starting weight showing no signs of disease. These data show that the YF/F17A double mutant is more attenuated than the individual NSP3 and N mutants (13, 14, 30). Overall, these results demonstrate that loss of SG antagonism by both NSP3 and N completely attenuated virus-induced weight loss in the SARS-CoV-2 hamster model of infection. Next, we examined viral replication in the upper and lower respiratory tract following infection with either WT or YF/F17A mutant. We observed no differences in nasal wash viral titers between WT and YF/F17A at 2- and 4 DPI (**Fig. 5C**); these results were consistent with prior studies that also observed no differences in the upper airway with the individual NSP3 and N mutants(14, 30). In contrast, we observed a consistent attenuation of YF/F17A mutant in lung viral titers compared to WT (**Fig. 5D**). The initial 1-log reduction in YF/F17A at 2 DPI expanded to 2-logs at 4 DPI. This level of attenuation was more consistent than both YF and F17A mutants individually (14, 30). In all animals, infection was resolved by 7 DPI in nasal washes and lung tissues. Together, these results show that disrupting the NSP3-FXR1 and N-G3BP1/2 binding interactions in combination drastically attenuates disease and viral load in lungs.

**Figure 5:**
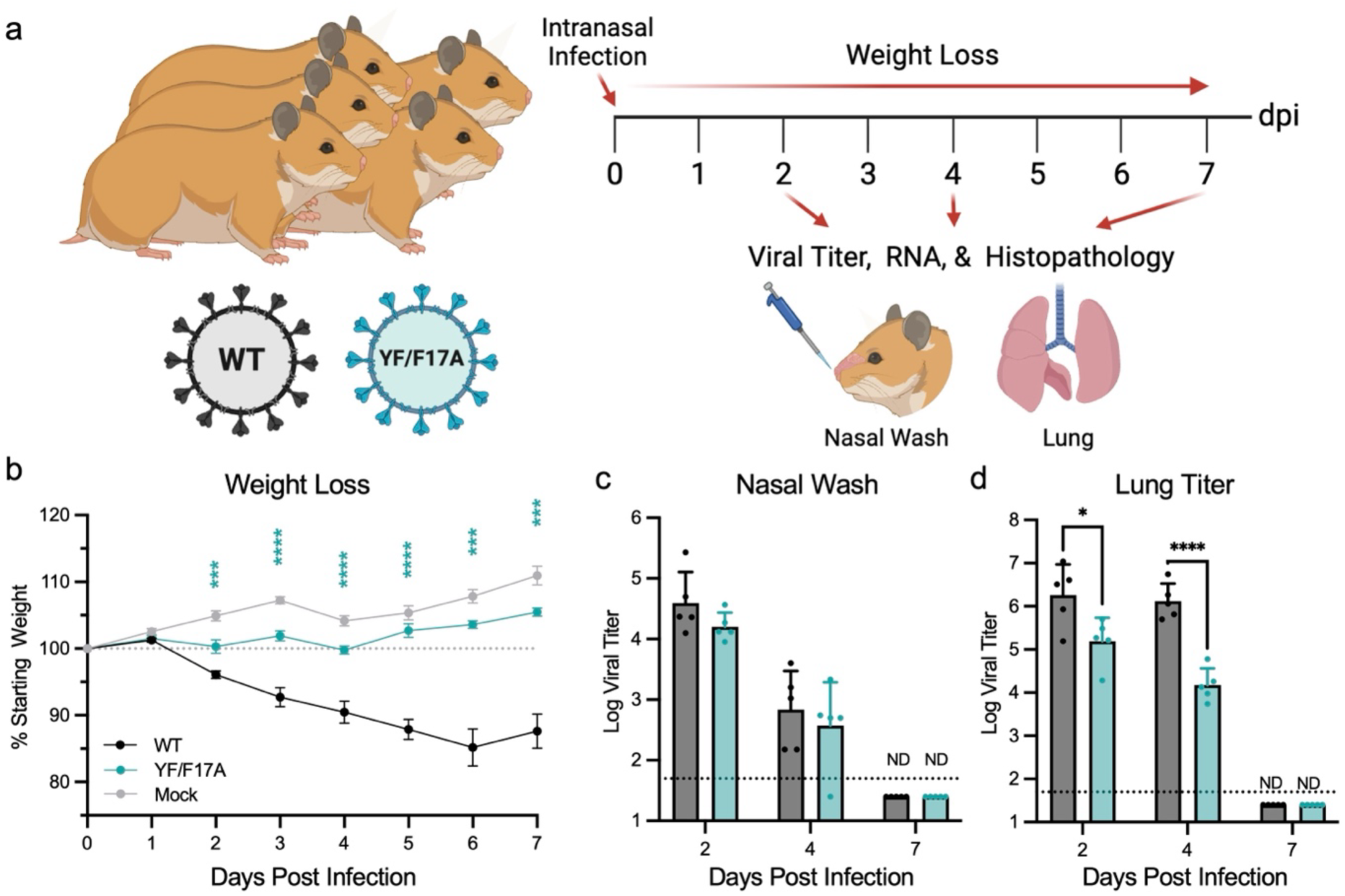
YF/F17A mutant has modest attenuation *in vivo*. (**a**) Schematic of golden Syrian hamster infection with WA-1 (WT) or YF/F17A mutant (teal) made using Biorender. (**b-d**) Three-to four-week-old male hamsters were intranasally inoculated with 10^5^ FFU of WT or YF and monitored daily for signs of disease and weight loss (**b**) over 7 days. Infectious titers were measured in the nasal wash (**C**) and lungs (**D**) on 2-,4-, and 7-DPI. Each dot represents an infected animal. The dotted lines represent the assay limit of detection. Statistical significance was determined by two-tailed Student’s t test. *P ≤ 0.05; **P ≤ 0.01; ***P ≤ 0.001, ****p<0.0001. Panel (a) was created with Biorender.com.

### YF/F17 mutant has reduced antigen staining in the lung

In addition to weight loss and replication, we also examined lung histopathology to evaluate changes between WT and the YF/F17A mutant. Staining for SARS-CoV-2 N revealed that viral antigen in the lung was reduced in the YF/F17A mutant relative to WT control (**S. Fig. 1A-E**). Beginning on day 2, we noted robust antigen staining in WT infected hamster lungs compared to the YF/F17A infected animals (**S. Fig. 1A-C**). The trend continued on day 4, with antigen largely absent in the YF/F17A group but maintained in the WT infected group (**S. Fig. 1C-E**). By day 7, viral antigen had been cleared from both WT and YF/F17A infected animals. While we noted no significant shifts between WT and YF/F17A mutant in antigen distribution between airway and parenchyma (**S. Fig. 1G-I**), these overall results indicate that less antigen in the lungs of YF/F17A infected hamsters corresponded with reduced lung viral titers **(Fig. 5D**).

### YF/F17A mutant has less lung immune infiltrate and inflammation

Examining immune infiltration and damage in the lung, we noted a significant attenuation in the YF/F17A mutant at late times post infection. Using Hematoxylin & Eosin (H&E) staining, a board-certified pathologist evaluated lung sections in a blinded manner from days 2, 4, and 7 following WT and YF/F17A infection (**S. Fig. 2A-F**). While differences were noted in viral antigen staining, WT and YF/F17A infected hamsters had largely similar immune infiltration at both days 2 and 4 post infection (**S. Fig. 2A-D**). For both WT and mutant infections, day 2 showed little evidence of immune infiltration compared to mock (**S. Fig. 2A-B, 2G**). By day 4, both infected groups showed increased immune infiltration and inflammation (**S. Fig. 2C-D**). However, day 7 had delineation between the infected groups with more severe immune infiltration in the WT infected hamsters (**S. Fig. 2E, 2H**). In contrast, immune infiltration and damage plateaued at day 4 level in the YF/F17A mutant and corresponded to reduced severity scores on day 7 (**S. Fig. 2F, 2H)** . Together, the H&E data indicates that the YF/F17A mutant produced reduced inflammation and immune infiltration in the lung. These data are consistent with reduced disease in terms of weight loss (**Fig. 5B**). Overall, these results show clear attenuation of the YF/F17A mutant in terms of *in vivo* disease which corresponded with the loss of SG control by the mutant virus.

### Host immune responses muted after YF/F17A mutant infection

We next evaluated the host responses in hamster lungs after infection with WT or YF/F17A. Total cellular RNA from infected lung at day 4 was sequenced with Poly(A)-ClickSeq (47, 48) (**Fig. 6**). At 4 DPI, we observed 3721 genes differentially expressed (P_adj_<0.1) between WT and YF/F17A. Tracing the fold change compared to mock (**Fig. 6A**), most host genes show correlation between WT and YF/F17A; however, some show anti-correlated expression patterns (up-regulated in WT vs. down-regulated in YF/F17A or vice versa). Examining gene ontology profiles, the YF/F17A mutant had reduced gene expression (P_adj_ < 0.1 and fold change < -2) of several host immune response pathways compared to WT (**Fig. 6B**). These pathways included genes associated with innate immune responses, inflammation, and cytokine/chemokine. In contrast, YF/F17A had augmented expression of genes associated with RNA polymerase II and other cellular factors, although the significance values of these pathways were lower than downregulation of immune pathways (**Fig. 6C**). Focusing further on the Go ontology, we examined RNA expression of genes related to innate immune responses: defense response to virus, immune response, inflammatory response, and innate immune response (**Fig. 6D**). For WT infection, in all 4 Go processes, we found upregulation of genes relative to mock infected hamsters (1^st^ column). While we observed some upregulation in the YF/F17A mutant relative to mock, these changes were muted or even inverse to what was observed in WT infected hamsters (2^nd^ column). Comparing mutant to WT directly, we observed that all the genes within these immune clusters had reduced expression in the YF/F17A mutant as compared to WT (3^rd^ column). The results indicated that the YF/F17A mutant has a muted immune response and suggest that SG activation during CoV infection limits the need for innate immune activation. Together, the data confirm prior experiments showing that an augmented innate immune response is not responsible for attenuation of the YF/F17A mutant.

**Figure 6:**
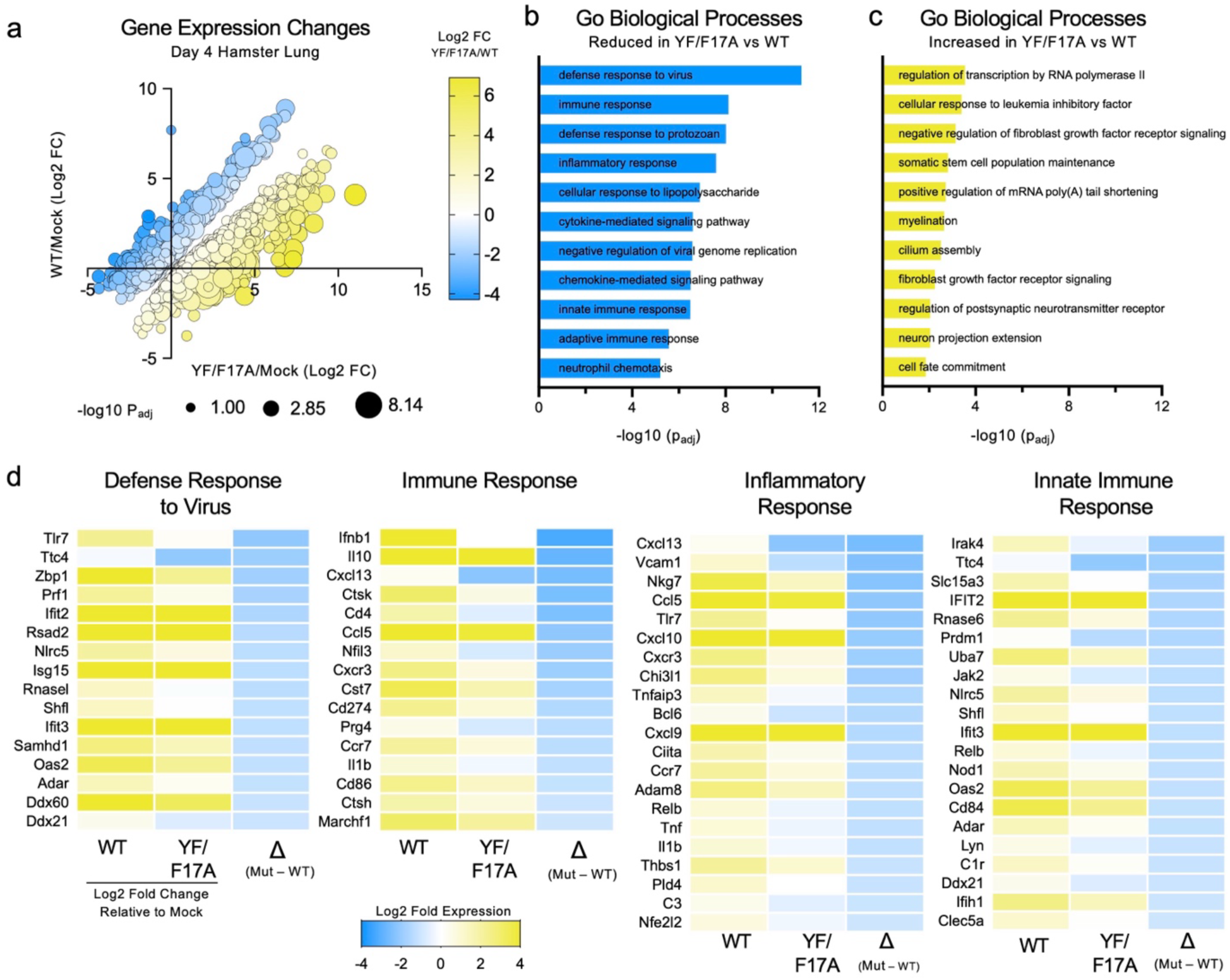
Host immune responses muted after YF/F17A mutant infection. (**a**) Host gene expression from day 4 hamster infected lungs comparing WT (Log2 fold changes vs mock, y-axis) to YF/F17A mutant (Log2FC vs mock) shows distinct patterns of up- and down-regulation. (**b-c)** Top gene ontology (GO) terms discovered from significantly (Padj < 0.1, |log2FC| > 1, N = 4 biological replicates) (**b**) down- or (**c**) up- regulated host genes (WT vs YF/F17A) at Day 4 post infection. (**d**) Day 4 gene expression analysis of individual genes in selected GO terms associated with immune responses to infection. Log2 Fold expression depicted for WT and YF/F17A mutant relative to mock as well as ratio of YF/F17A to WT expression.

### YF/F17A mutant infection protects against SARS-CoV-2 rechallenge

The loss of NSP3 and N SG antagonism renders SARS-CoV-2 significantly attenuated both *in vitro* and *in vivo*. Notably, the YF/F17A mutant not only has reduced viral load but also diminished innate immune responses. This fact distinguishes it from other SARS-CoV-2 mutants that are inherently replication deficient or induce exacerbated immune responses (49). It also leaves the possibility that reduced immune responses disrupt adaptive immunity, impacting protection from subsequent challenge. To explore this question, we determined in the YF/F17A mutant could provide protection against subsequent rechallenge with WT SARS-CoV-2. Briefly, golden Syrian hamsters were initially challenged with 10^5^ FFU SARS-CoV-2 WT, YF/F17A mutant, or mock. The infected animals were monitored for weight loss and disease over a seven-day time course and had similar kinetics as initial studies (**S**. **Fig. 3**). After 21 DPI, we rechallenged all groups with 10^5^ FFU of WT SARS-CoV-2 virus and monitored daily for weight loss and clinical signs of disease (**Figure 7A**). At days 2, 4, and 7 post rechallenge, serum, nasal wash, and lung specimens were collected for analysis (**Fig. 7A**). As expected, mock vaccinated hamsters had significant weight loss following challenge with WT SARS-CoV-2 (**Fig. 7B**); these animals lost weight beginning at day 2 and we observed peak weight loss at day 6. In contrast, hamsters previously vaccinated with WT SARS-CoV-2 or the YF/F17A mutant had no weight loss following infection. The results indicate that despite reduced immune activation, infection with YF/F17A mutant could protect from disease following rechallenge with WT SARS-CoV-2.

**Figure 7:**
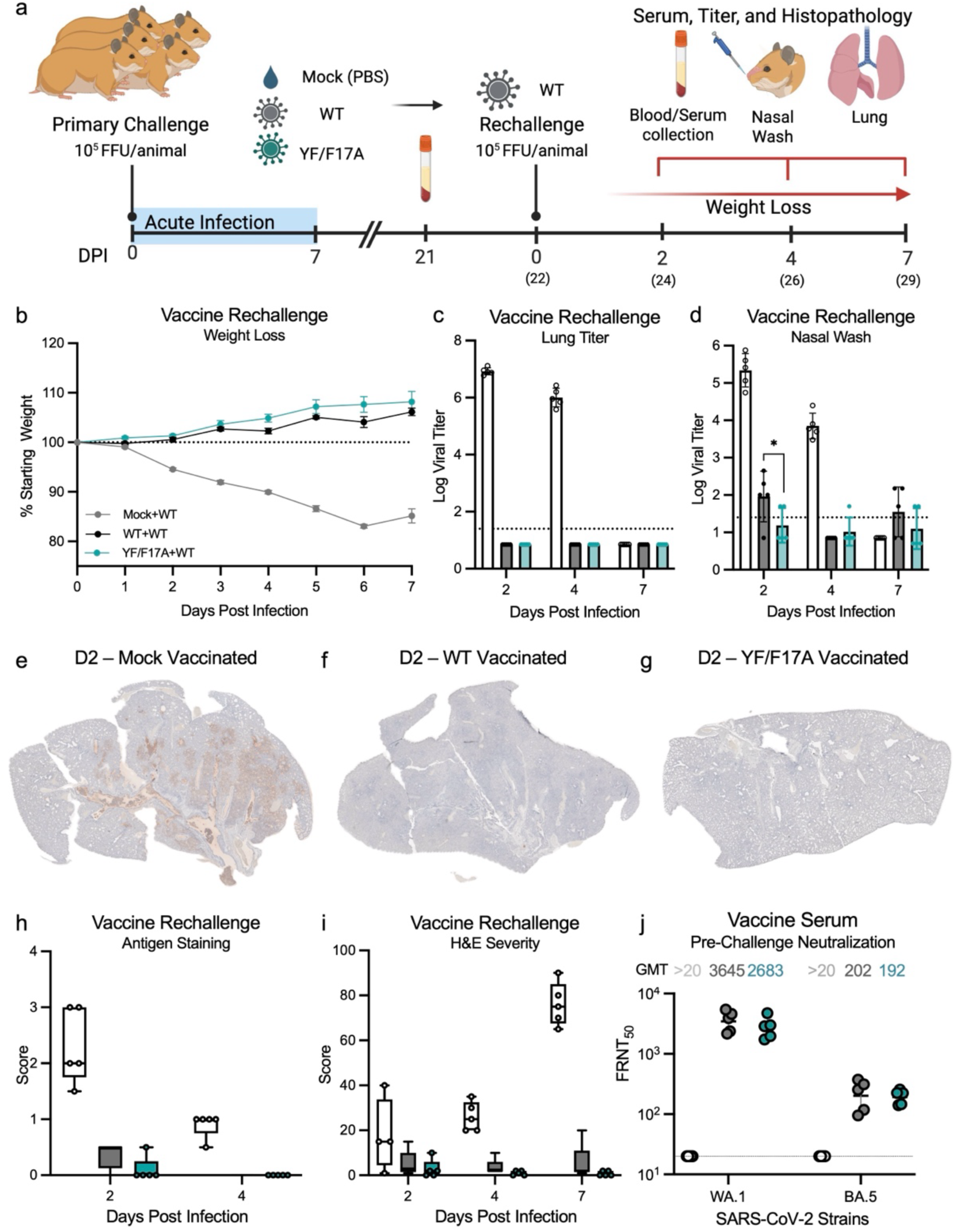
Y**F/F17A infection protects against rechallenge with SARS-CoV-2 WT. (a**) Schematic of vaccination and rechallenge experiments. For primary challenge, 3-to 4-week-old male hamsters were intranasally vaccinated with 10^5^ FFU of WT (grey), YF/F17A mutant (teal), or mock (PBS, white), and monitored daily for disease and weight loss. At 21 days post-primary infection, hamsters were bled to collect serum and rechallenged the next day (22 DPI/D0) with 10^5^ FFU of WT. (**b-d**) Rechallenged animals were monitored daily for weight loss (**b**). Infectious titers were measured in the lung (**c**) and nasal wash (**d**) on days 2, 4, and 7 post rechallenge. (**e-g)** Nucleocapsid antigen staining of left lung section from (**e**) mock-vaccinated, (**f**) WT-vaccinated, or (**g**)YF/F17A mutant vaccinated hamsters 2 days post rechallenge. (**h)** Antigen staining was scored in a blinded manner by for lung total score for mock vaccinated (white), WT-vaccinated (grey) or YF/F17A-vaccinated (teal) infected sections at days 2 and 4 post rechallenge. (**i**) H&E staining was scored in a blinded manner for total score for mock vaccinated (white), WT-vaccinated (grey) or YF/F17A-vaccinated (teal) infected sections at days 2, 4,7 post rechallenge. (**j**) Neutralizing antibody responses measured by live-virus based focus reduction neutralization test (FRNT50) assays against SARS-CoV-2 WA1 and BA.5 using pre-challenge sera from mock-vaccinated (white), WT-vaccinated (grey), and YF/F17A vaccinated (teal). FRNT50 geometric mean titers (GMTs) are indicated above each group. Each dot represents experimental value from an infected animal. The dotted lines represent the assay limit of detection. Statistical significance was determined by two-tailed Student’s t test. *P ≤ 0.05; **P ≤ 0.01; ***P ≤ 0.001, ****p<0.0001. Figure 4.7A was created with Biorender.com.

We subsequently measured the viral load in the upper and lower respiratory tracts of animals. Consistent with earlier studies (**Fig. 5**), mock vaccinated animals had robust viral replication in both the lung and nasal wash after infection (**Fig. 7C & D**). Examining lung titer, the mock vaccinated animals had significant viral loads at both day 2 and day 4 post infection (**Fig. 7C**). In contrast, both WT and YF/F17A vaccinated animals had no detectable virus at either time point. Examining nasal wash, the mock infected animals had robust titers at both day 2 and 4 post infection (**Fig. 7D**). While titers were significantly diminished in the WT and YF/F17A mutant nasal wash samples, detectable virus was observed in both groups. These results are consistent with incomplete protection provided by infection/vaccination in the upper airway compared to control (50). Notably, we did observe a small, but significant difference in day 2 nasal wash as the WT had slightly higher load than YF/F17A mutant. Overall, the results indicate that despite the reduced immune induction during primary infection, the YF/F17A mutant limits viral replication to similar levels as vaccination with the WT SARS-CoV-2 strain.

We subsequently examined lung histopathology from control and vaccinated hamsters (**Fig. 7E-I, S. Fig. 4**). Utilizing antigen staining for N, we found that mock vaccinated hamsters had significant staining in both the airways and parenchyma following infection at both days 2 and 4 (**Fig. 7E**). In contrast, both WT and YF/F17A vaccinated animals showed minimal antigen staining in lung section consistent with lung viral titer data (**Fig. 7F-H**). Examining immune pathology by H&E staining, we again observed robust inflammation and immune infiltrate in the mock vaccinated hamsters infected with WT SARS-CoV-2 that progressed over the 7-day time course (**Fig. 7I**, **S**. **Fig. 4A-C**). In contrast, no significant immune infiltrate or inflammation was observed in the WT and YF/F17A group following rechallenge with WT SARS-CoV-2 (**Fig. 7I, S**. **Fig. 4D-I)**. Together, the results indicate that protection against both viral replication and disease was achieved by YF/F17A mutant vaccination at level similar to WT SARS-CoV-2 vaccination.

Finally, we examined serum neutralization from hamsters vaccinated with WT or YF/F17A mutant, or mock. Briefly, sera were collected from vaccinated and unvaccinated hamster groups 21 days post vaccination and prior to rechallenge with SARS-CoV-2 WT (**Fig. 7A**). FRNT50 assays were performed as previously described (51) and demonstrated that both WT and YF/F17A mutant vaccination had similar neutralization of SARS-CoV-2 strains (**Fig. 7J**). Examining the backbone SARS-CoV-2 WA.1 strain, WT and YF/F17A mutant neutralizations were similar in their log geometric mean titer (GMT). Similarly, while neutralization against SARS-CoV-2 Omicron variant BA.5 had a 10-fold reduction compared to WA.1, both WT and YF/F17A vaccination resulted in nearly identical values. Together, the results indicate that the muted immune responses following primary infection with YF/F17A mutant did not significantly impair vaccination efficacy. Overall, the serum findings combined with the challenge results indicate targeting SG antagonism has utility as a mechanism for live attenuated vaccines for SARS-CoV-2 and potentially other CoVs.

## Discussion

SARS-CoV-2 devotes two viral proteins, NSP3 and N, to antagonize SG activity suggesting a critical need to block this function. The YF/F17A mutant confirmed this reality rendering the virus unable to replicate efficiently in cells (**Fig. 1**) or cause significant disease in hamsters (**Fig. 5, S. Fig. 1-2**). Importantly, the YF/F17A mutant attenuation was not associated with augmented innate immune responses. Cell culture studies indicated that innate immune responses were not augmented in the YF/F17A mutant (**Fig. 2**) and hamster lung RNA showed reduced activation of immune pathways (**Fig. 6**). Instead, attenuation of the YF/F17A mutant correlates with more activated SGs as measured by G3BP1 foci in infected cells (**Fig. 3 - 4**). Over time, this activation corresponded to fewer infected cells indicating that SGs are likely responsible for the deficit in the YF/F17A mutant (**Fig. 1 & 5**). Overall, these results show the YF/F17A mutant is more attenuated than the individual YF and F17A mutants alone. In addition, the data highlight the importance of disruption of SGs in SARS-CoV-2 infection and pathogenesis.

Both N and NSP3 are multifunctional proteins in the context of coronavirus infection with a variety of domains and roles during infection. During SARS-CoV-2 infection, the interplay between the two viral proteins allows kinetic control of SG formation. At early times, NSP3 uses its YF motif in its highly variable region to bind FMRP proteins and disrupt SG induction (14, 30). While less potent than N, NSP3 serves to delay SGs while N accumulates and performs other roles during the early parts of CoV infection. At late times (10 HPI), N functions to bind to G3BP1/2, master regulators of SGs, functionally disabling their activity during the latter part of infection (13, 16). In the absence of both SG antagonists, SARS-CoV-2 has attenuated replication and reduced pathogenesis that corresponds to accumulation of SGs in infected cells. Yet, SGs may only partially be responsible for the limitations of the YF/F17A mutant. Studies have indicated that while N binds to G3BP1 to disrupt SGs, it may also facilitate key aspects of viral replication via trafficking G3BP1 to the viral replication complex and aiding viral mRNA translation (18, 52). In this context, the hijacking of G3BP1 may have dual role of being both pro-viral and antagonizing host SG responses. Coupled with the loss of early disruption by NSP3, the YF/F17A mutant is more limited by SG activation than each of the individual mutants (13, 14).

In addition to defining a critical need to control this pathway, our study also provides insights toward the mechanism of SG mediated attenuation of SARS-CoV-2. The interplay between SGs and host innate immunity have led to mixed results in recent years. Some groups have found that SGs activation amplifies innate immune responses (53–55). While other studies have argued that SGs modulate innate immune activation by acting as shock absorbers for overt activation (56). Still, other groups have reported no link between SGs and innate immune signaling (57, 58). While we do not explore this question in depth, our results indicate that loss of SG antagonism does not induce an augmented innate immune response to SARS-CoV-2 in cells (**Fig. 2**). Similarly, we find that interferon and inflammatory responses are reduced following *in vivo* infection with the YF/F17A mutant (**Fig. 6**). These results suggest SG activation may reduce innate immune activation, although attenuated YF/F17A mutant replication may also signal changes due to reduced viral load. Given these caveats, our results indicate that in the context of SARS-CoV-2 infection, loss of SG antagonism does not amplify innate immune responses.

Unlinking SG activity and innate immunity has potential implications for treating and preventing SARS-CoV-2 infection. Approaches to disrupt CoV infection for vaccination typically rely on reducing viral replication by targeting a key viral process or activating host immune responses via interferon pathways (59). Here, disrupting antagonism of SGs allows control of the SARS-CoV-2 infection through a distinct attenuation pathway independent of type I IFN. Using the YF/F17A mutant as a live attenuated vaccine, we found that animals infected with the mutant had minimal disease during acute infection (**Fig. 5)**. Subsequent rechallenge with WT-SARS-CoV-2 demonstrated protection from both viral replication and disease at level similar to WT vaccination (**Fig. 7**). Despite the reduced immune activation, the YF/F17A mutant produced similar neutralizing titers as WT and coupled with robust protection from disease, indicated feasibility of this approach for live attenuated vaccination. Importantly, since the molecular mechanisms governing SG antagonism are known for SARS-CoV-2, a combination approach targeting both NSP3 interaction with FMRP proteins and N interactions with G3BP1 may have utility as a drug treatment. In fact, studies have targeting the N-G3BP1 with drugs have shown some initial efficacy against SARS-CoV-2 (60, 61). Pairing this inhibition with NSP3/FMRP disruption may offer a novel, effective therapeutic pathway to activate SGs and inhibit SARS-CoV-2 infection.

Moving beyond SARS-CoV-2, these findings have implications for understanding the role of SGs in other coronaviruses. For N, examination of the N-terminal domain indicate conservation of F17 in similar locations of other CoVs including SARS-CoV implying the N-G3BP1 function may be broadly conserved within the Sarbecovirus family. In contrast, while the HVR residues are broadly conserved in the Sarbecovirus family, a similar motif is not found in the other CoV families (30). Importantly, other CoV families have developed other methods to disrupt SG functions directly. For example, MERS-CoV has been implicated in disrupting SG responses with mutants lacking the ORF4a viral protein inducing more SGs (62). More broadly, CoVs employ several conserved viral proteins that disrupt double-stranded RNA production or immune recognition including N, NSP1, NSP15, and NSP14 (13, 25, 27, 41, 63–70). With dsRNA acting as stimulator of SG activation, disruptions of these viral antagonists have also been linked to SG activation during CoV infection (71–73). Together, these findings show that CoVs must spend significant genetic capital to modify and shape SG activation. As such, targeted approaches to disrupt this antagonism and activating SGs may be a viable approach for treatments and vaccine approaches across the entire CoV family.

Our findings also have implications for studying SGs in the context of viral infection. Many viruses utilize viral proteins to disrupt, delay, or inhibit SG functions including influenza, alphaviruses, herpes viruses and others. For decades, groups have utilized these viral families to explore the mechanisms and factors that contribute to SG activation and function. Among them, influenza NS1 has been consistently identified as a key SG antagonist and numerous studies have utilized influenza NS1 mutant to identify key host factors that shape SG responses (74). Notably, influenza NS1 has several roles in viral infection including antagonism of SG response and innate immunity (75). These overlapping roles and functions make interpretations of results more difficult as numerous aspects of influenzas infection in addition to SG activity are impacted. Here, we show that the YF/F17A mutant of SARS-CoV-2 maintains viral replication, lacks induction of innate pathways, and induces a significant increase in SGs contrasting to WT virus. With the recent downregulation to a lower biocontainment level, these results make the SARS-CoV-2 YF/F17A mutant a potential useful tool to study and understand SG activity in the context of immunity to viral infection.

Overall, our study demonstrates the importance of controlling SG activity to SARS-CoV-2 infection and pathogenesis. Loss of NSP3 and N mechanisms to block SG formation results in significant attenuation of the SARS-CoV-2 mutant *in vitro* and *in vivo*. This attenuation corresponds to significant SG activation in the mutant, and notably, similar to reduced innate immune activation as compared to WT SARS-CoV-2 following infection. The result is reduced viral replication and disease *in vivo*. Overall, the results highlight the importance of both N and NSP3 antagonism and highlight a novel pathway targeting SG antagonism as a means for CoV attenuation and treatment.

## Materials & Methods

### Cell Lines

African green monkey kidney Vero E6 cells were cultured in high glucose Dulbecco’s modified Eagle medium (DMEM) (Gibco #11965-092) supplemented with 10% Fetal Clone II (FBS) (Hyclone; #SH30071.03) and 1% antibiotic/antimycotic (anti-anti) (Gibco #5240062). Vero E6 expressing human TMPRSS2 were grown in high glucose DMEM supplemented with 10% FBS, 1% anti-anti, and 1 mg/mL Geneticin (G418) (Gibco; 10131035). Calu-3-2b4 (Calu-3) cells were grown in high glucose DMEM with 10% defined FBS, 1% anti-anti, and 1 mg/mL sodium pyruvate (1mM). A549 cells expressing human ACE2 were grown in 10% FBS, 1% anti-anti, and 10 µg/mL Blasticidin S HCL (Gibco; A1113902).

Vero E6 and Calu-3 cells were provided by Dr. Ralph Baric. A549-ACE2 cells were provided by Dr. Chien-Te Kent Tseng. All cell lines grow at 37°C with 5% CO_2_ and are mycoplasma tested periodically. (Last negative test in 2024). Single cell FACS sorting (BD Aria Fusion) was applied on Vero-TMPRSS2 and A549-ACE2 cells to select for TMPRSS2 or ACE2 positive cell populations.

### Viruses

***Reverse Genetic Clones***. All recombinant viruses were generated using reverse genetics as previously described (31, 32). WT and mutant SARS-CoV-2 sequences are used from the USA-WA1/2020 isolate sequence provided by the Word Reference Center for Emerging Viruses and Arboviruses (WRCEVA), originally obtained from the US Centers for Disease Control and Prevention (CDC) as previously described (34). The NSP3 YF mutant was constructed using standard cloning techniques as previously outlined for our reverse genetics system (31, 32) with virus stocks being amplified on Vero E6-TMPRSS2 to prevent mutations from occurring in furin cleavage site (FCS) 33482650. All viruses were expanded once (P1) for use in studies and titered by Focus-forming assay (FFA) and plaque assay (for plaque morphology). All infections and manipulations were performed in a biosafety level 3 (BSL-3) laboratory in accordance with approved protocols and use of appropriate personal protective equipment. P1 viral stocks were verified through Sanger sequencing of cDNA for mutations in spike furin cleavage site, the NSP3-HVR, and N-F17A for the introduced mutations.

### Biosafety

The synthetic construction of SARS-CoV-2 NSP3 mutants was reviewed for DURC/P3CO policies and approved by the University of Texas Medical Branch Institutional Biosafety Committee. All studies in animals were conducted under a protocol approved by the UTMB Institutional Animal Care and Use Committee and complied with USDA guidelines in a laboratory accredited by the Association for Assessment and Accreditation of Laboratory Animal Care. UTMB is a registered Research Facility under the Animal Welfare Act. It has a current assurance (A3314-01) with the Office of Laboratory Animal Welfare (OLAW), in compliance with NIH Policy. Procedures involving infectious SARS-CoV-2 were performed in the Galveston National Laboratory ABSL3 facility.

### In vitro infection

***Viral replication kinetics*.** Viral replication studies were performed as previously described (30). Briefly, cells (Vero E6, A549-ACE2) were seeded in seeded in six well plates the day before infection. Calu-3 cells were seeded at high density and allowed to growth to confluency in six wells plates before conducting growth curve. Cells were infected at indicated MOI for 45 minutes at 37°C with 5% CO_2_ and every 15 minutes. After absorption, plates were washed three times with PBS, and fresh medium was replaced in wells, indicating time zero. Wells were sampled at specified time points. Three technical replicates were collected for each timepoint, and each experiment was performed twice. Samples were titrated with focus-forming assay.

For high MOI infections, A549-ACE2 cells were plated in 12-well plates at 5×10^5^ cells per well. Viruses were diluted in PBS and added to cells and incubated for 45 minutes at 37°C with 5% CO_2._ and rocked every 15 minutes. Cells were washed three times and replaced with complete DMEM (10% FBS with 1x Anti-Anti). For each time point, 200 ul of supernatant was collected for viral titers, and cells were lysed for qPCR gene expression analysis with Trizol ™.

### Type I IFN Assay

For IFN pretreatment, 100 Units (U)/mL of recombinant IFN-β (PBL Laboratories) was added to Vero E6 cells 16-18 hours prior to infection. Growth curves were conducted as mentioned above.

### Real-Time quantitative PCR (qPCR)

***RNA Extraction & qPCR*.** Total RNA from cells were isolated with Direct-zol RNA Miniprep Plus kit (Zymo, #R2072) according to manufacturer’s protocol. RNA was reverse transcribed to complementary DNA (cDNA) using the iScript cDNA synthesis kit according to manufacturer’s instructions (Bio-Rad, 1708891). RT-qPCR was performed on cDNA using the Luna Universal qPCR Master Mix (NEB #M3003) per manufacturer’s instructions. Fluorescent readings were measured on a Bio-Rad CFX Connect instrument using Bio-Rad CFX Maestro 1.1 (version 4.1.2433.1219). The primers used to amplify host targets were human GAPDH (forward:5’-TCAAGATCATCAGCAATGCC ; reverse: 5’- AAGTTGTCATGGATGACCTTGG), IFIT1 (forward:5’-CCAAGGAGACCCCAGAAACC ; reverse: 5’-CGCTACGTGGAGTGAGCTAG), IFIT3 (forward:5’-AAGAACAAATCAGCCTGGTCAC ; reverse: 5’-TCCCTTGAGACACTGTCTTCC), IFNβ (forward:5’-AGTAGGCGACACTGTTCGTG ; reverse: 5’-AGCCTCCCATTCAATTGCCA), OAS1 (forward:5’-GAGCTCCTGACGGTCTATGC ; reverse: 5’-TCATCGTCTGCACTGTTGCT), MDA5 (forward:5’-AAGCCCACCATCTGATTGGAG ; reverse: 5’-CCACTGTGGTAGCGATAAGCAG),PKR (forward:5’-GAAGTGGACCTCTACGCTTTGG ; reverse: 5’- GATGATGCCATCCCGTAGGTC) .Average Ct values of technical replicates were normalized using the ΔΔCt method to the house keeping gene (human GAPDH) and expressed as Log_2_ relative fold change.

### Immunofluorescence and microscopy

A549-ACE2 cells were seeded on 96-well black sided plates one day prior to infection. Infections were conducted at indicated MOI in an inoculum volume of 50 µL volume per well. Incubation was conducted as mentioned in viral replication kinetics. Wells were replaced with fresh cell culture medium containing no selective antibiotics. To induce SG formation, wells were replaced with cell culture media containing 1mM sodium arsenite 30 minutes prior to inactivation timepoint. Plates were inactivated in 10% buffered formalin at indicated HPI. Antibody staining was adapted from Lamichhane et. al, 2024. Briefly, fixed cells were permeabilized with 0.2% Triton X-100 in PBS for 10 minutes at rt and blocked with 2% BSA for 1 hour. Samples were then incubated with primary antibodies; Rabbit anti-G3BP1 (Cell signaling #61559) and mouse anti-Nucleocapsid (Cell Signaling, 68344BC) overnight at 4◦C. The next day, cells were washed 3 times with PBS and incubated with host-specific Alexa Fluro 488 (Invitrogen; A32731) and Alexa Fluro 555 (Invitrogen; A28180) secondary antibodies for 1 hour at rt, away from light 39221199.^45^ After secondary incubation, plates were washed 3 times, and nuclei were stained with 1 ug/ml Hoecsht 3342 (Thermo Fisher, H1399) according to manufacturers instructions for 15 minutes at rt. Images were captured using a Cytation 7 instrument at 20x magnification.

### Stress granule quantification

Quantification of SGs was adapted from Dolliver et. Al., 2022 36534661. At least four areas were imaged in one well (repeated in triplicate) with consistent exposure setting for all images. Infected (N-positive) cells containing at least 1 granule we counted as SG+ using the multi-point tool in ImageJ 36534661.^46^ The total number of cells expressing N was also counted using the multi-point tool in ImageJ. Percentage stress granule positive was then calculated by dividing the resulting number. For robust image analysis, standardized image acquisition practices were followed. Low to moderate cell densities were imaged to prevent negative impacts on cell counting and segmentation. Image acquisition was carried out using optimal constant exposure settings for each experiment. The “QuickFigures” plugin for ImageJ was used to prepare immunofluorescence panels (76).

### *In vivo* Infection

***Animal studies.*** Three-to-four-week male Syrian golden hamsters (HsdHAN:AURA strain) were purchased from Envigo, Indianapolis, IN. For infection, hamsters were intranasally inoculated with 1×10^5^ FFU of SARS-CoV-2 WT or YF/F17A mutant in a 100 µL volume. Infected animals were weighed and monitored daily for illness over 7 days. Cohorts of 5 animals per group (including mock infected animals) were anesthetized with isoflurane, and nasal washes were collected in 400 µl PBS on endpoint days (2, 4, and 7 DPI) for titers. Animals were then humanely euthanized by CO_2_ immediately following nasal washes for lung specimen collection. Lobes were stored in RNAlater solution (Invitrogen) for qPCR analysis and in PBS for titer analysis.

***Rechallenge study***. For rechallenge experiments, hamsters were intranasally inoculated with 1×10^5^ FFU of SARS-CoV-2 WT or YF/F17A mutant in a 100 µL volume PBS. Infected animals were weighed and monitored daily for illness over 7 days. Mock (PBS) challenged groups were given 100 µL volume PBS. At day 21 post-primary challenge, serum was collected from all animals via retroorbital bleeding. Animals were allowed to recover and challenged at day 22 post-primary challenge. Infected animals were weighed and monitored daily for illness over 7 days. Specimen collections were performed as previously described, with the addition of serum collection via cardiac puncture.

### Focus-forming assay

Focus-forming assays (FFAs) were performed as previously described with adaptations (77). Briefly, Vero E6 cells were seeded in 96-well black walled plates to be 100% confluent the next day. Cell culture supernatants, nasal washes, or homogenized tissues containing SARS-CoV-2 underwent 10-fold serial dilutions in serum-free media. Twenty µl of diluted and undilute sample were added to wells and incubated for 45 minutes with rocking every 15 minutes. After absorption, methylcellulose overlay (0.85%) was added to each well, and cells were incubated at 37°C for 24 hours. The next day, methylcellulose was removed, and cells were washed three times with PBS and fixed in 10% buffered formalin for 30 minutes at room temperature (rt) to inactivate virus. For staining, cells were permeabilized and blocked with PBS solution containing 0.1% saponin and 0.1% BSA (Perm. Buffer) for 30 minutes. Following incubation, α-SARS-CoV-2 Nucleocapsid primary antibody (Cell Signaling, 68344) at 1:3,000 in perm. buffer overnight at 4°C. Cells were washed 3 times with 1X DPBS before incubating with Alexa Fluor 555-conjugated α-mouse secondary antibody (Invitrogen, A28180) at 1:2,000 in Perm. Buffer for 1 hour at rt wrapped in foil. Cells were washed 3 times with 1X DPBS, and fluorescent foci images were captured using a Cytation 7 cell imaging multi-mode reader (Biotek). Foci were counted using ImageJ free software (NIH).

***Homogenization***. Hamster lungs were infected as described in ‘Animal Studies’ and the right inferior lobes were stored in sterile PBS, zirconia beads, and kept at -80◦C. Tissues were homogenized using a MagNALyser (Roche Life Science) at 6,000 rpm for 1 minute total.

### Histology

Left lung lobes were harvested from hamsters and fixed in 10% buffered formalin solution for at least 7 days. Fixed tissue was then embedded in paraffin, cut into 5 µM sections, and stained with hematoxylin and eosin (H&E) on a SAKURA VIP6 processor by the University of Texas Medical Branch Surgical Pathology Laboratory.

### Immunohistochemistry

Fixed and paraffin-embedded left lung lobes from hamsters were cut into 5 µM sections and mounted onto slides by the University of Texas Medical Branch Surgical Pathology Laboratory. Paraffin-embedded sections were warmed at 56°C for 10 min, deparaffinized with xylene (3x 5-min washes) and graded ethanol (3x 100% 5-min washes, 1x 95% 5-min wash), and rehydrated in distilled water. After rehydration, antigen retrieval was performed by steaming slides in antigen retrieval solution (10 mM sodium citrate, 0.05% Tween-20, pH 6) for 40 min (boil antigen retrieval solution in microwave, add slides to boiling solution, and incubate in steamer). After cooling and rinsing in distilled water, endogenous peroxidases were quenched by incubating slides in TBS with 0.3% H_2_O_2_ for 15 min followed by 2x 5-min washes in 0.05% TBST. Sections were blocked with 10% normal goat serum in BSA diluent (1% BSA in 0.05% TBST) for 30 min at room temperature. Sections were incubated with primary anti-N antibody (Sino #40143-R001) at 1:1000 in BSA diluent overnight at 4°C (13, 78). Following overnight primary antibody incubation, sections were washed 3x for 5 min in TBST. Sections were incubated in secondary HRP-conjugated anti-rabbit antibody (Cell Signaling Technology #7074) at 1:200 in BSA diluent for 1 hour at room temperature. Following secondary antibody incubation, sections were washed 3x for 5 min in TBST. To visualize antigen, sections were incubated in ImmPACT NovaRED (Vector Laboratories #SK-4805) for 3 min at room temperature before rinsed with TBST to stop the reaction followed by 1x 5-min wash in distilled water. Sections were incubated in hematoxylin for 5 min at room temperature to counterstain before rinsing in water to stop the reaction. Sections were dehydrated by incubating in the previous xylene and graded ethanol baths in reverse order before mounted with coverslips.

### Next Generation Sequencing (NGS) libraries

For NGS analyses, RNA template was extracted from animal tissue with Direct-zol RNA miniprep kits (Zymo Research). To understand the transcriptomic changes of infected animals, extracted total cellular RNA from hamster lungs were subjected to PolyA-ClickSeq (47), which utilizes a semi-anchored oligo(dT) primer (5’- (T)_21_-3’) to specifically target polyA tails of cellular mRNA and a 1:5 azido-ddNTP:dNTP ratio to ensure sufficient termination of cDNA. The PolyA-ClickSeq library was constructed with previously established protocol (79) and gel selected libraries were single-end sequenced with ElementBio Aviti.

### Bioinformatics and data analysis

Raw reads sequenced from PolyA-ClickSeq libraries are processed and analyzed with previously established *DPAC* (80) pipeline (https://github.com/andrewrouth/DPAC/) with parameters “*-p PMBCD*”, which detects and processes the polyA-containing reads and maps the upstream sequence of polyA tail to host reference. In this study, a PolyA-site clustering database was curated based on the published reference genome and annotation of *Mesocricetus auratus* (Genbank accession number: GCA_000349665.1). Due to the incompleteness of the reference genome, the genome mapping criterion was slightly loosened in hisat2 (81) stage with parameter *“--score-min L,0,-0.3*”. Differential gene expression profiles of mock (PBS), WT, and YF/F17A infected hamster lung tissues were then analyzed with DESeq2 (82) that has been integrated in the *DPAC* pipeline (D stage) to reveal the normalized read count of each annotated gene.

For differential gene expression analyses, hierarchical gene clustering was conducted with *Cluster 3.0* (http://bonsai.hgc.jp/~mdehoon/software/cluster/). This is followed by TreeView (http://jtreeview.sourceforge.net/) to visualize the heat map and gene clusters. A gene is considered differentially expressed with Benjamini and Hochberg *P*_adj_ < 0.1 and |log2FC| > 0.585. Gene ontology assay was conducted with DAVID (83) and differentially expressed genes (*P*_adj_ < 0.1, |log2FC| > 1), to highlight the most direct GO terms in biological process, cellular component, and molecular function. All plots and other statistics are conducted on GraphPad Prism (v.10.6.1).

### Focus reduction neutralization test

FRNT assays were performed as previously described (51, 84–86). Briefly, serum samples in duplicate were threefold diluted in eight serial dilutions using Dulbecco’s modified Eagle’s medium with an initial dilution of 1:10. Serially diluted samples were mixed with an equal volume of the desired SARS-CoV-2 variant (100 to 200 foci per well). The virus-serum mixtures were incubated at 37°C for 1 hour in a round bottom 96-well culture plate. After 1 hour of incubation, the virus-serum mixture was added to VeroE6-TMPRSS2 cells and incubated at 37°C for an additional hour. Post-incubation, the mixture was removed from cells, and 100 μl of prewarmed 0.85% methylcellulose overlay was added to each well. Plates were incubated at 37°C for 18 to 40 hours (depending on variants). After the appropriate incubation time, the methylcellulose overlay was removed, and cells were washed with phosphate-buffered saline (PBS) and fixed with 2% paraformaldehyde for 30 min. After fixation, cells were washed twice with PBS and permeabilized using permeabilization buffer for at least 20 min. After permeabilization, cells were incubated with an anti–SARS-CoV-2 spike antigen primary antibody directly conjugated to Alexa Fluor 647 (CR3022-AF647) overnight at 4°C. Cells were then washed twice with 1× PBS and imaged on an ELISPOT reader (CTL Analyzer).

## Data Availability

The raw data that support the findings of this study are available from the corresponding author upon reasonable request. The raw sequencing data have been deposited with links to NCBI BioProject databased under PRJNA1477634.

## Acknowledgements

Research was supported by grants from NIAID of the United States NIH (R01-AI153602, R21-AI145400, U19-AI171413 to VDM; R21-AI183054 to BAJ). The research was also supported by STARs Award provided by the University of Texas System and Burroughs Welcome Fund Investigators in Pathogenesis grant to VDM. Further funding was provided by the American Lung Association (1413648). Trainee funding provided by NIAID of the NIH to REA (T32AI007526-24). We thank Parimal, Meagan Rippee-Brooks, Alyssa M McLeland, Adam Hage for technical assistance on the manuscript.

## Author Contributions

Conceptualization: REA, MSS, BAJ, VDM

Formal analysis: REA, JC, KGL, DHW, BAJ, VDM Funding acquisition: REA, BAJ, VDM

Investigation: REA, JC, KGL, YZ, LKE, ALM, YA, XD, LL, AM, WM, JAP, DHW, VDM

Methodology: REA, JC, KGL, XX, BAJ, VDM Project Administration: KSP, MSS, XX, BAJ, VDM Supervision: KSP, XX, MSS, BAJ, VDM Visualization: REA, JC, VDM

Writing – original draft: REA, VDM

Writing – review and editing: REA, JC, DHW, BAJ, VDM

---

**S. Figure 1:**
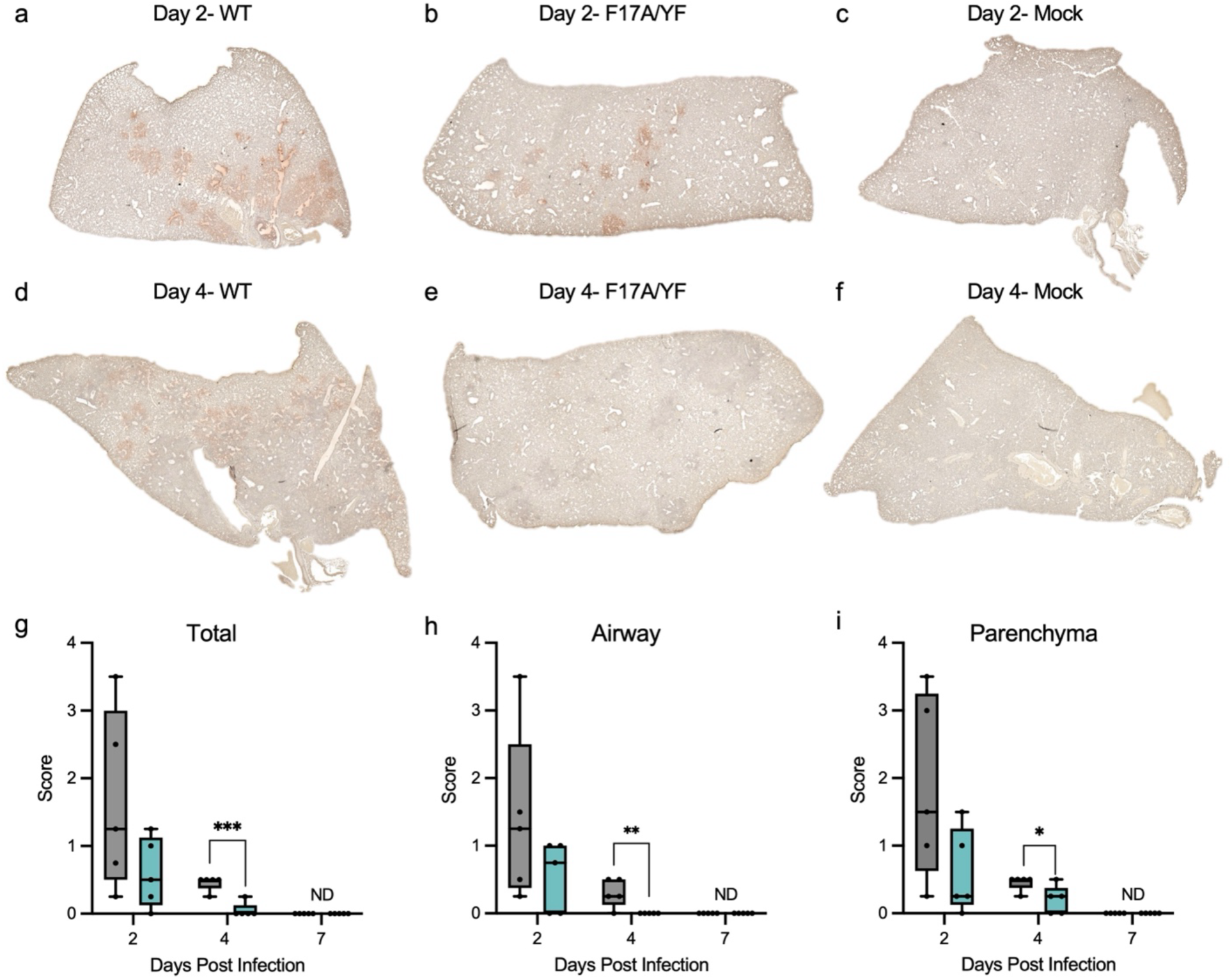
Viral antigen staining confirms YF/F17A attenuation *in vivo*. (**A-F**) Nucleocapsid antigen staining of left lung section from hamsters infected with 10^5^ FFU of either WT, YF/F17A mutant or mock PBS at (**A-C)** 2 or (**D-F**) 4 DPI. Antigen staining was scored in a blinded manner by location in the (**G**) lung total, (**H**) parenchyma, and (**I**) airway for WT (grey) or YF/F17A (teal) infected lungs. Statistical analysis was conducted using a two-tailed Student’s *t* test. \**P* ≤ 0.05; \*\**P* ≤ 0.01; and \*\*\**P* ≤ 0.001.

**S. Figure 2:**
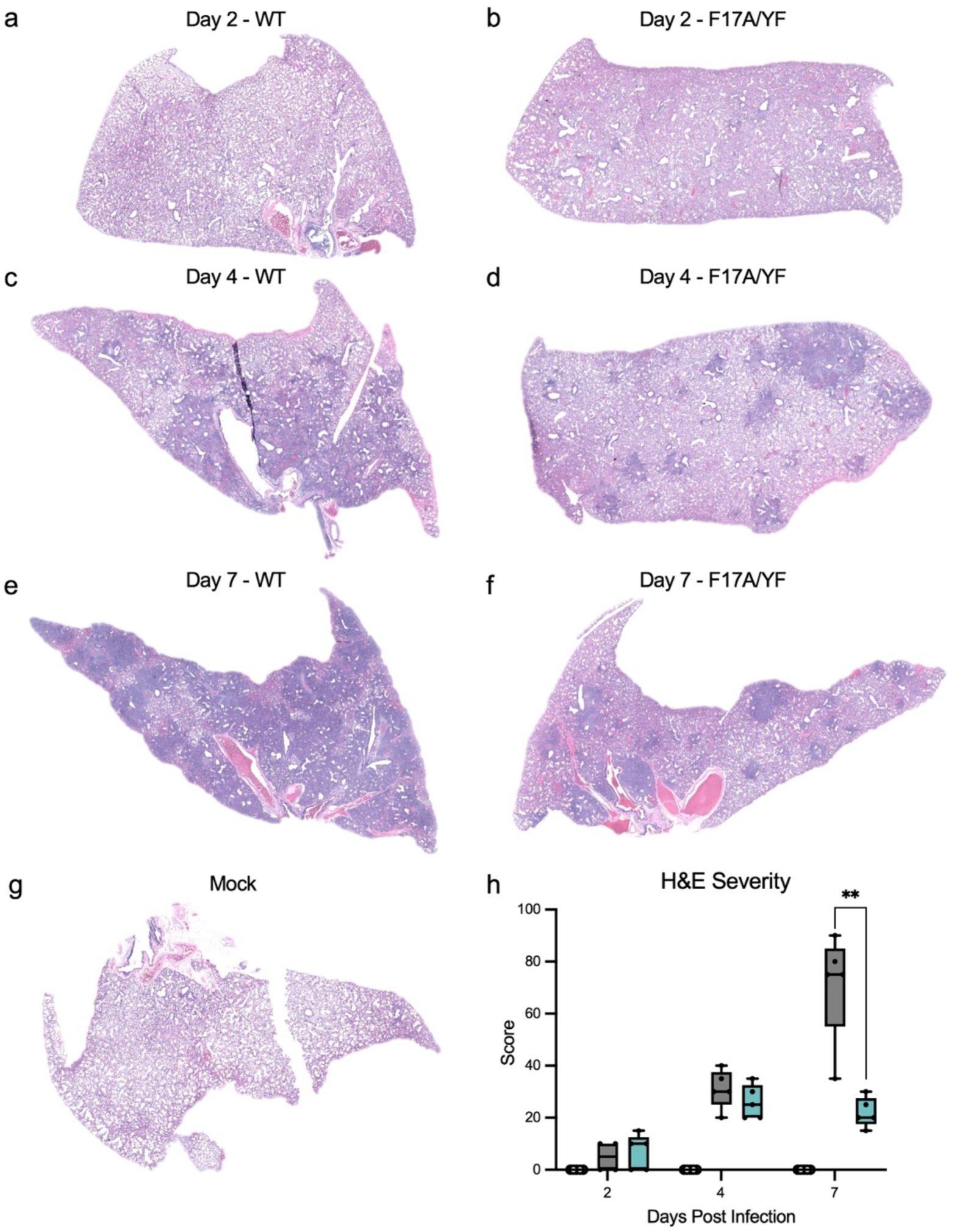
YF/F17A infection reveals reduced immune infiltration and damage. Reduced inflammation and damage in YF/F17A infected lungs. (**A-G**) Representative H&E staining of the left lung of hamsters infected with 10^5^ FFU of either WT or YF/F17A SARS-CoV-2 at (**A-B**) 2-, (**C-D**) 4-, or (**E-F**) 7 DPI or (**G**) mock. (**H**) Mock, WT (black), or YF/F17A (teal) lung sections from each day were scored for histopathological analysis, with sections from individual animals averaged and representing a single point. Statistical analysis was conducted using a two-tailed Student’s t test. *P ≤ 0.05; **P ≤ 0.01; ***P ≤ 0.001.

**S. Figure 3:**
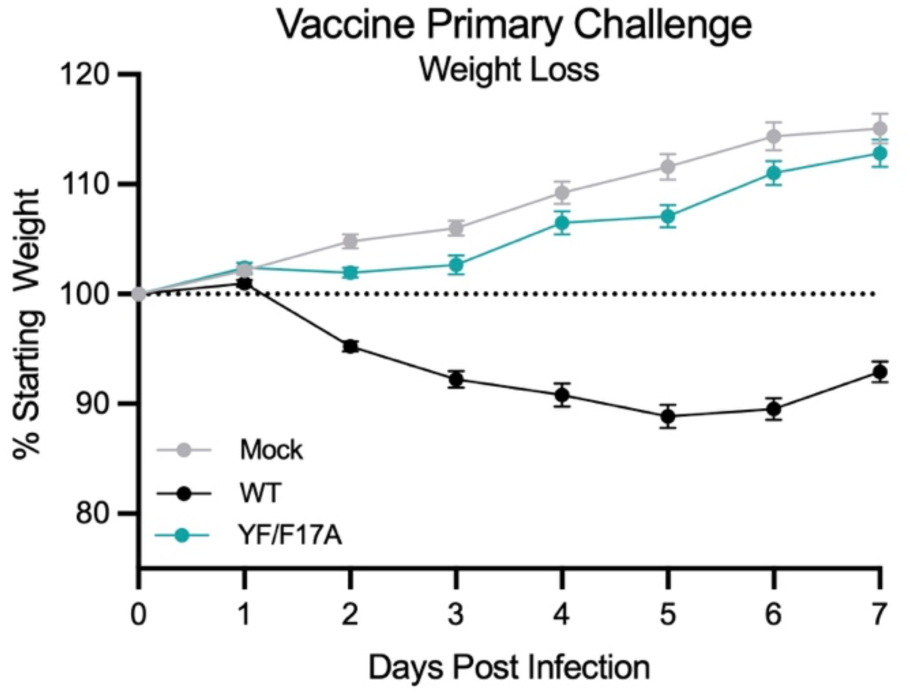
YF/F17A infection causes similar disease in vaccination as primary infection. For vaccination, three-to four-week-old male hamsters were intranasally inoculated with 10^5^ FFU of WT, YF/F17A mutant, or equal volume PBS (mock), and monitored daily for signs of disease and weight loss over a 7 day time course.

**S. Figure 4:**
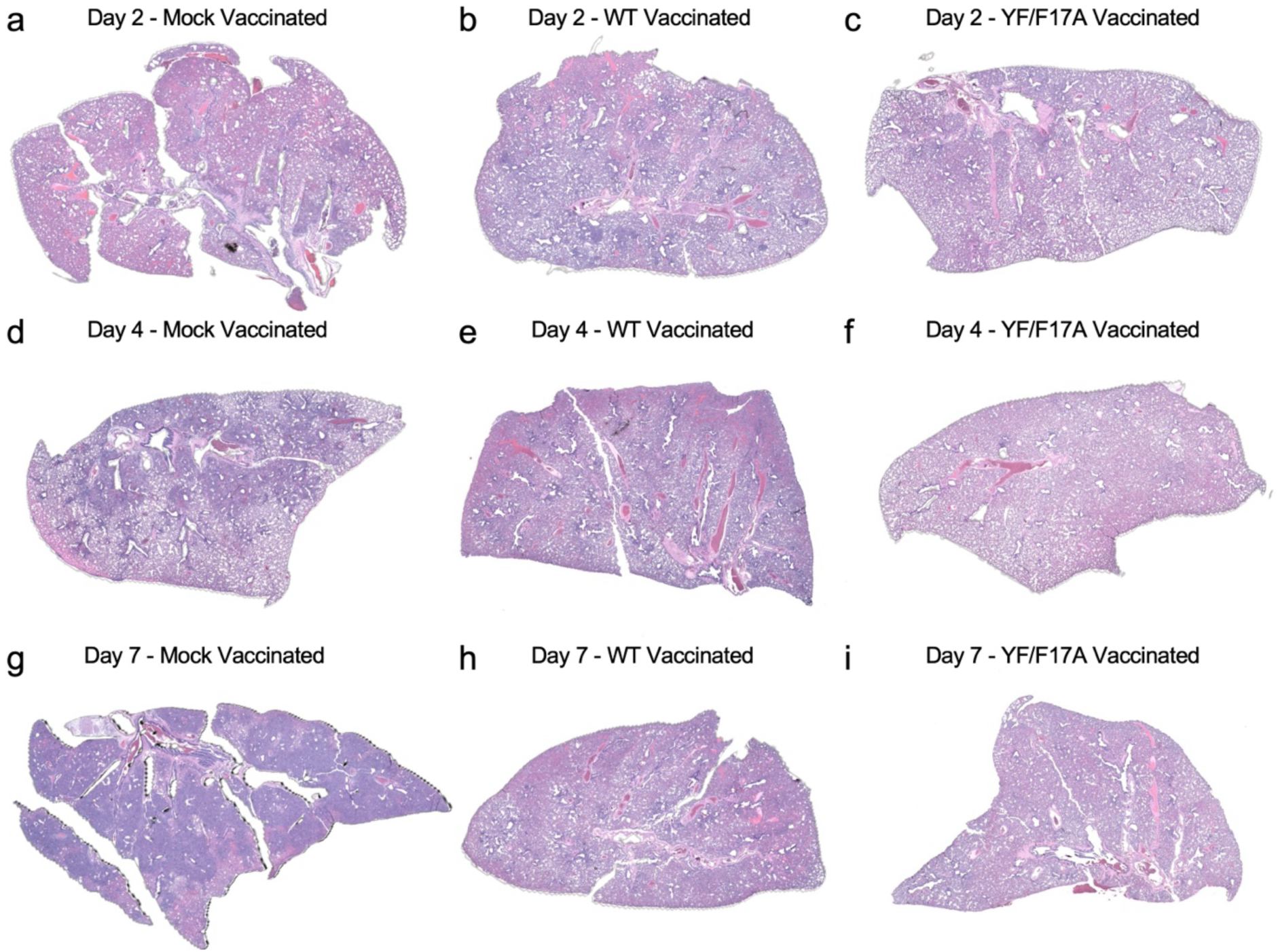
YF/F17A infection reveals reduced immune infiltration and damage. Minimal inflammation or damage observed in vaccinated WT and YF/F17A infected hamster lungs. (**A-G**) Representative H&E staining of the left lung section of hamsters from mock vaccinated, WT-vaccinated or YF-F17A vaccinated rechallenged with 10^5^ FFU of WT SARS-CoV-2 at (**A-C)** 2-, (**D-F**) 4-, or (**G-I**) 7 DPI or (G) mock.

